# Letter to the Editor regarding “Long-read genome sequencing provides novel insights into the harmful algal bloom species *Prymnesium parvum*” by Jian et al. (2024)

**DOI:** 10.64898/2026.01.21.699772

**Authors:** Jennifer H. Wisecaver, Timilehin Jeje, Nathan F. Watervoort

## Abstract

Jian et al. (2024) describe *de novo* genome assemblies for two strains of *Prymnesium parvum* (sensu lato, s.l.), a cryptic species complex of toxic, unicellular algae responsible for harmful algal blooms around the world. Here, we present evidence that the labels for UTEX 2797 and CCMP 3037 were inadvertently swapped by Jian et al. (2024). This resulted in sequence data labeled “UTEX 2797” but derived from strain CCMP 3037, and vice versa. Strain misidentification is a major risk with cryptic species like *P. parvum* s.l., and our reanalysis of the data in Jian et al. (2024) underscores the urgent need for clade-specific markers to ensure accurate and efficient strain identification.

---

*Prymnesium parvum* s.l. is a toxic, bloom-forming haptophyte composed of three main clades referred to as A, B, and C which produce different forms of the toxin prymnesin (Binzer et al. 2019). Recently, two publications generated reference assemblies for the strains UTEX 2797 and CCMP 3037 (Table 1) (Wisecaver et al. 2023; Jian et al. 2024). UTEX 2797 and CCMP 3037 are both members of clade A, characterized by the production of prymnesin type A (Figure 1). Wisecaver et al. (2023) further subdivided clade A into three subclades. In their analysis, the A1 subclade was comprised of two strains from the United States: 12B1 isolated from Moss Creek Lake, Texas in 2010 and CCMP 3037 isolated from Twin Buttes Lake, Wyoming in 2008. The A2 subclade consisted of two strains from Europe: RCC 3703 isolated in 1953 from Millport in the United Kingdom and CCMP 2941 isolated in 2007 from Lake Repnoye, Russia. The last A subclade proposed by Wisecaver et al (2023) consisted of two strains: UTEX 2797 and 12A1 both isolated from Texas waterbodies in 2002 and 2010, respectively. Both UTEX 2797 and 12A1 showed evidence of hybridization between the A1 and A2 subclades. Specifically, both strains had elevated levels of heterozygosity compared to the other 13 strains in the analysis. Both strains also contained roughly twice as much DNA content per nucleus (∼280 Mbp), as estimated via flow cytometry, than their k-mer-based estimate of genome size (∼120 Mbp). Wisecaver et al (2023) assembled the genome of UTEX 2797 using a combination of Oxford Nanopore Technologies (ONT) long reads, Illumina short reads, and Hi-C reads for scaffolding. They performed a phylogenomic analysis of the genes on the resulting 66 scaffolds and showed that most scaffolds formed syntenic pairs with genes on one member of the scaffold-pair grouping with the A1 subclade and genes on the other member of the scaffold-pair grouping with the A2 subclade. In a breadth of coverage analysis, 12A1 Illumina reads mapped well to all UTEX 2797 scaffolds while other type A strains mapped to only one scaffold per syntenic pair;

**Table 1.**
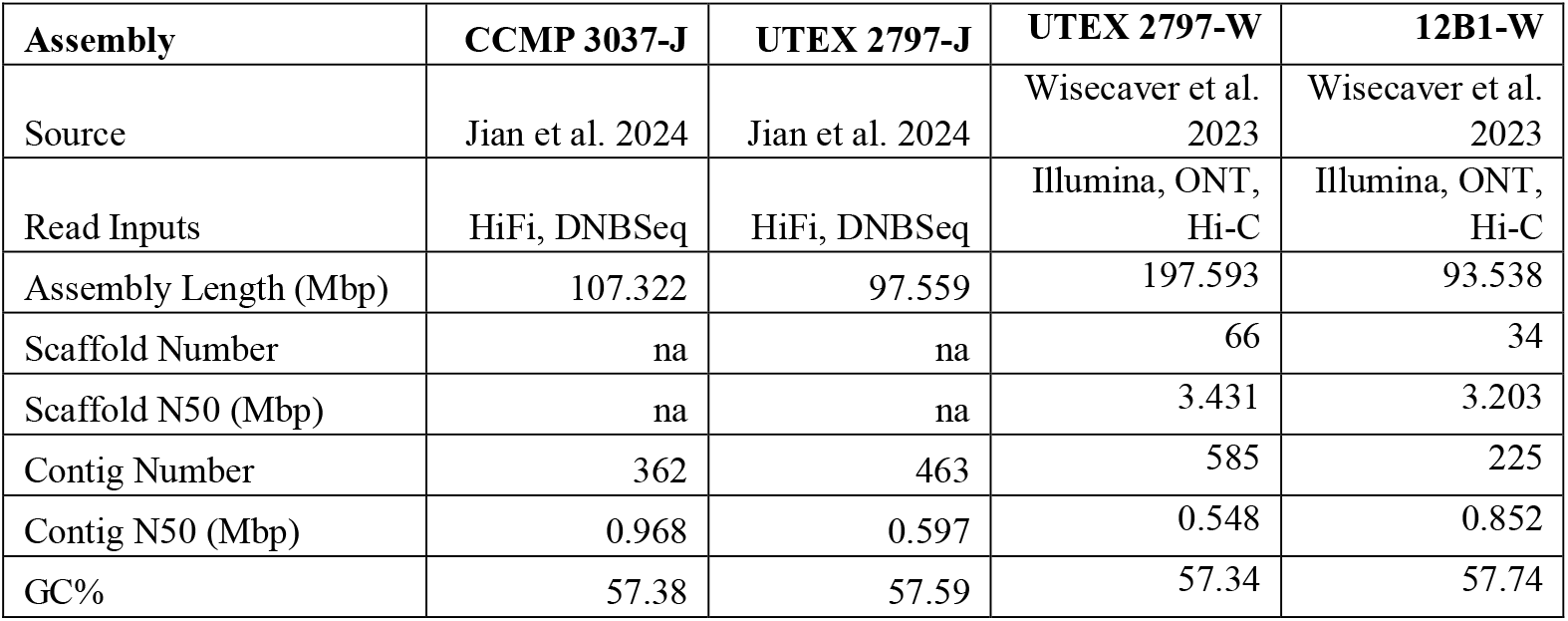
Statistics from all A-type *P. parvum* assemblies.

**Figure 1.**
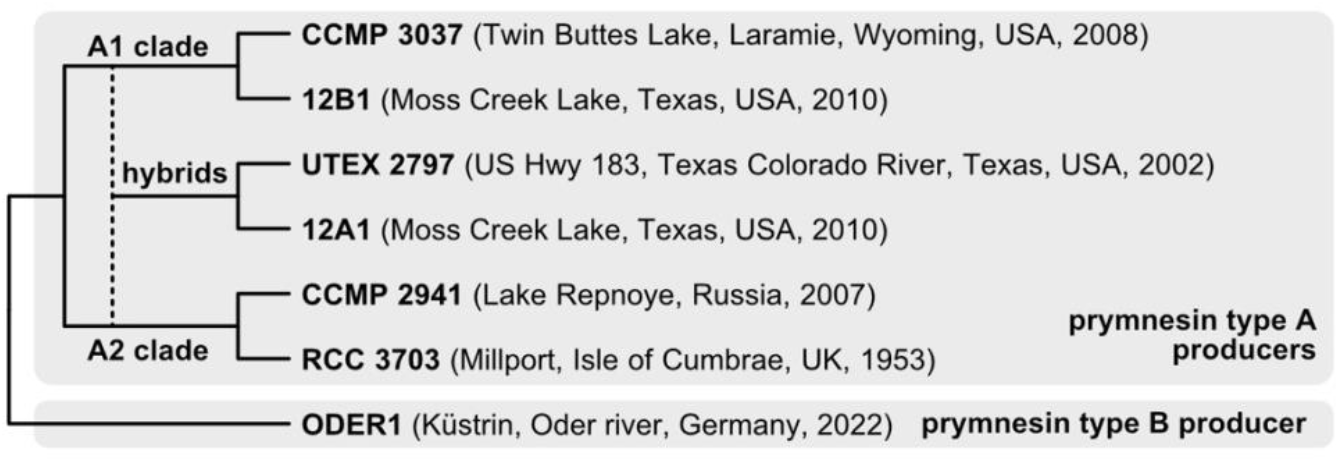
*P. parvum* s.l. phylogeny. Model of evolutionary relationships between *P. parvum* s.l. strains present in this analysis based on phylogenetic and genomic analyses performed by Wisecaver et al. (2023).

A1 strains mapped to the A1-like scaffolds in UTEX 2797, and A2 strains mapped to the A2-like scaffolds. Taken together, the data suggested that UTEX 2797 and 12A1 were formed via hybridization between A1-like and A2-like parents with the apparent retention of the full, or near full, genomic compliment of each parent (Figure 1).

The UTEX 2797 and CCMP 3037 assemblies produced by Jian et al. (2024) disagreed with the analysis by Wisecaver et al. (2023) and instead placed UTEX 2797 in the A1 clade. In addition to the obvious issues resulting from having contradicting assemblies in the field, UTEX 2797 is a commonly used laboratory strain featured in many publications (e.g., Blossom et al. 2014; Rashel and Patiño 2017; Lysgaard et al. 2018; Binzer et al. 2019; Richardson and Patiño 2021; Chávez Montes et al. 2024). The existing confusion surrounding UTEX 2797 has led to its exclusion from recent analyses of the *P. parvum* s.l. complex (e.g. Kuhl et al. 2024). To correct the record surrounding a strain important to the field of *P. parvum* research and to highlight the difficulties of analyses within cryptic species complexes, we subjected nucleic acid sequences from UTEX 2797 and CCMP 3037 to a battery of tests. This includes newly generated sequences for this analysis and sequences from third-party research groups.

The k-mer plots for the CCMP 3037 and UTEX 2797 libraries generated by Wisecaver et al. (2023), Jian et al. (2024), and this study were compared to each other. In the Jian et al. (2024) data, strain CCMP 3037 is highly heterozygous, as indicated by the two distinct peaks in the k-mer frequency plot (Figure 2A). Conversely, strain UTEX 2797 shows only one dominant k-mer peak, indicative of far less heterozygosity (Figure 2B). The pattern is the opposite in the Wisecaver et al. (2023) data: CCMP 3037 had low heterozygosity (Figure 2C), while UTEX 2797 was high (Figure 2D). Re-sequenced reads demonstrate the same pattern of heterozygosity as previously published by Wisecaver et al. (2023) and the opposite pattern of what was published by Jian et al. (2024) (Figure 2E,F). This potentially indicated that the strain IDs were switched for the Jian et al. (2024) investigation.

**Figure 2.**
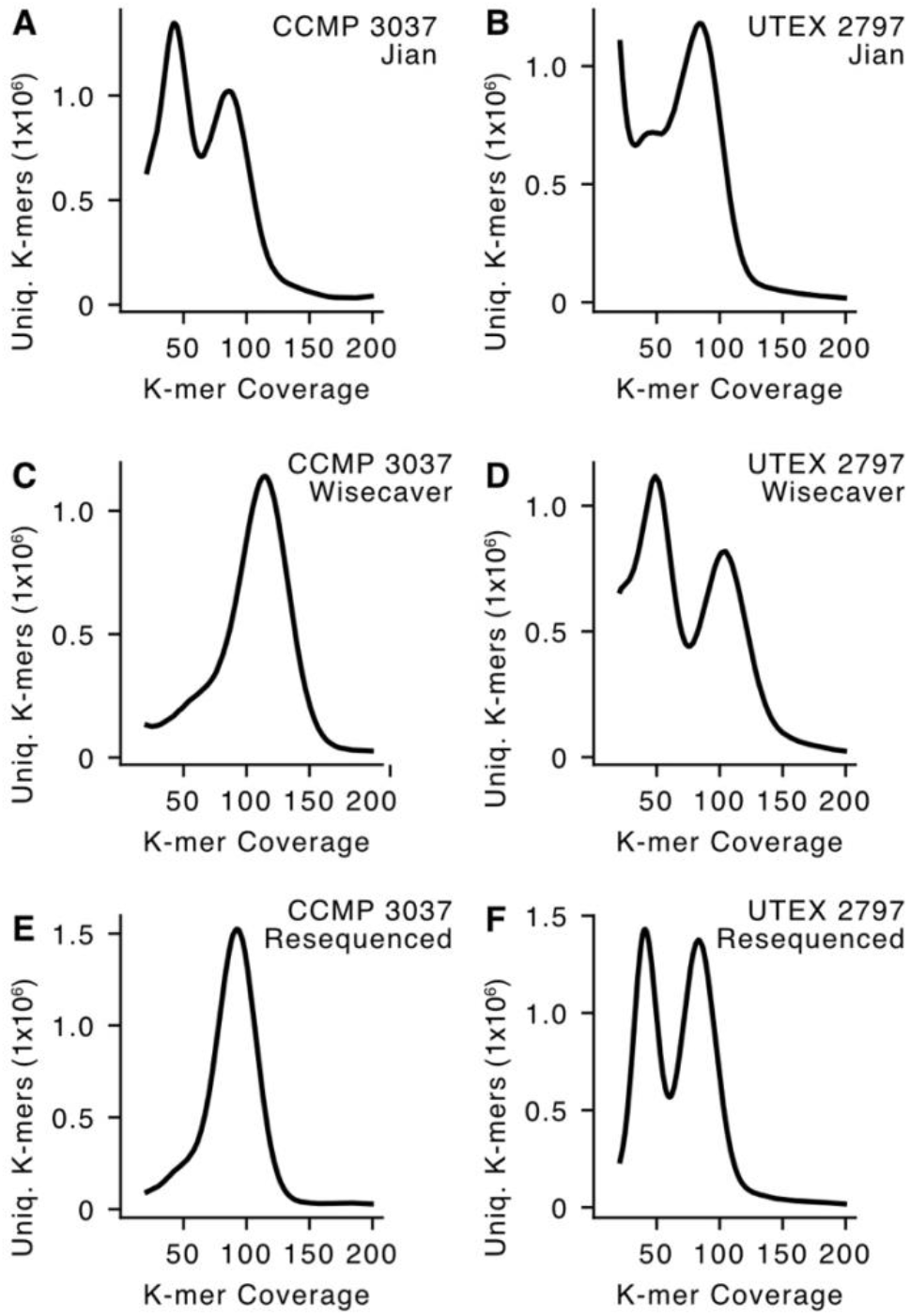
K-mer frequency plots for CCMP 3037 and UTEX 2797. Data from Jian et al. (A,B); from Wisecaver et al. (C,D); and resequenced for this study (E,F).

In a principal component (PC) analysis of the resulting SNPs, the strains clustered into three distinct groups along the first PC axis (Figure 3A). These three clades generally correspond to the phylogenetic clades proposed by Wisecaver et al. (2023). The first group contains strains from the A2 subclade, CCMP 2941 and RCC 3703. The second group contains strains of the A1 subclade, 12B1 and the CCMP 3037, sequenced by Wisecaver et al. (2023). The Jian et al. UTEX 2797 and the freshly reacquired and resequenced CCMP 3037 also cluster in this first group. The Illumina reads from Wisecaver CCMP 3037, Jian UTEX 2797, and the resequenced CCMP 3037 almost entirely overlap one another in PC space, which illustrates the strong similarity between the three samples and indicates that they correspond to the same strain. The third group contains the two hybrid strains proposed by Wisecaver et al. (2023), 12A1 and UTEX 2797. The CCMP3037 from Jian et al. (2024) and the freshly reacquired and resequenced UTEX 2797 also cluster within this third group. As in the first cluster, the Illumina reads from Wisecaver UTEX 2797, Jian CCMP 3037, and resequenced UTEX 2797 overlap one another. Critically, sequences from a third party also confirm our assessment. The UTEX 2797 RNAseq reads independently sequenced by Kuhl et al. (2024) also cluster in this third group demonstrating that our original UTEX 2797 (Wisecaver et al., 2023), the freshly reacquired and resequenced UTEX 2797 (this letter), the UTEX 2797 used for RNAseq (Kuhl et al., 2024) and the CCMP 3037 in Jian et al. (2023) are all UTEX 2797.

**Figure 3.**
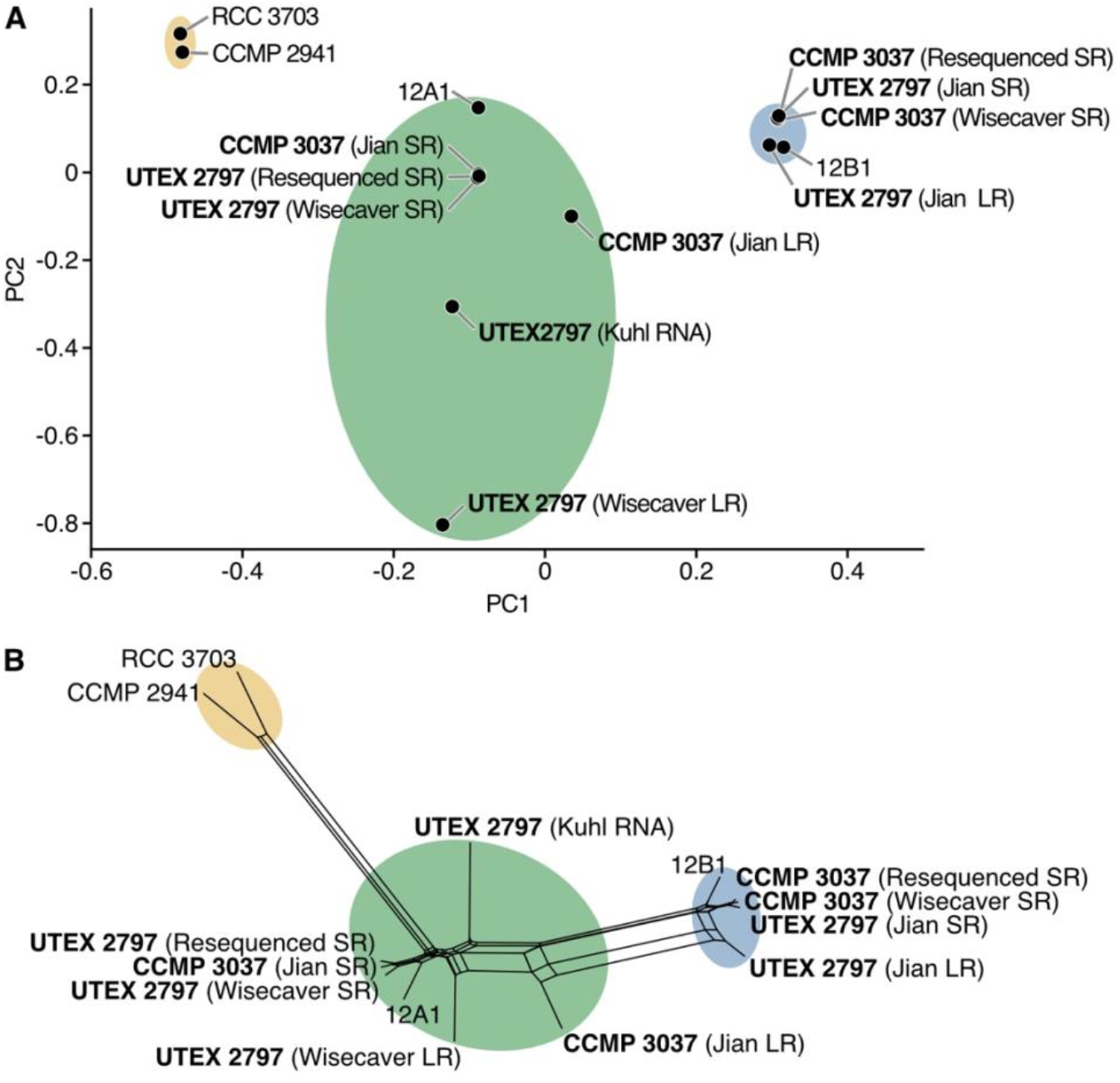
SNP analysis of *P. parvum* s.l. type A strains. A) PCA analysis B) Phylogenetic network. LR, long read; SR, short read.

All filtered variants were used to create a NeighborNet phylogenetic network (Huson 1998; Bryant 2003), which recovered the same trends observed in the PCA (Figure 3B). UTEX 2797 from Jian et al (2024) grouped with the A1 strains from Wisecaver et al (2023), and CCMP 3037 from Jian et al (2024) grouped with the hybrid strains from Wisecaver et al (2023). UTEX 2797 from Kuhl et al (2024) also grouped with the hybrid strains from Wisecaver et al. (Figure 3B). Taken together, these analyses show that the CCMP 3037 and UTEX 2797 strain labels were swapped in Jian et al. (2024).

Lastly, we generated k-mer frequency plots for the long-read libraries of CCMP 3037 and UTEX 2797 from Jian et al. (2024). These plots were dominated by k-mers with low coverage (< 5), and the plots showed no visible coverage peaks (data not shown). We then noted that the long-read libraries provided through China National GeneBank DataBase (CNGBdb, accession number CNP0004284) have significantly fewer reads and a lower read depth than those reported by Jian et al. (2024) (Table 2). The deposited long reads are sufficient for approximately 6.5x coverage for both strains. Hifiasm (Cheng et al. 2021), the assembler used by Jian et al. (2024), recommends at least quadruple this coverage in its FAQ (∼13x per haplotype; for diploids = 26x total coverage). We do not know the reason for the discrepancy between the quantity of reads deposited and the numbers reported in the original manuscript. However, N50 is significantly higher in the deposited dataset (Table 2), indicating that the difference could be due to filtration of short reads prior to deposition.

**Table 2.**
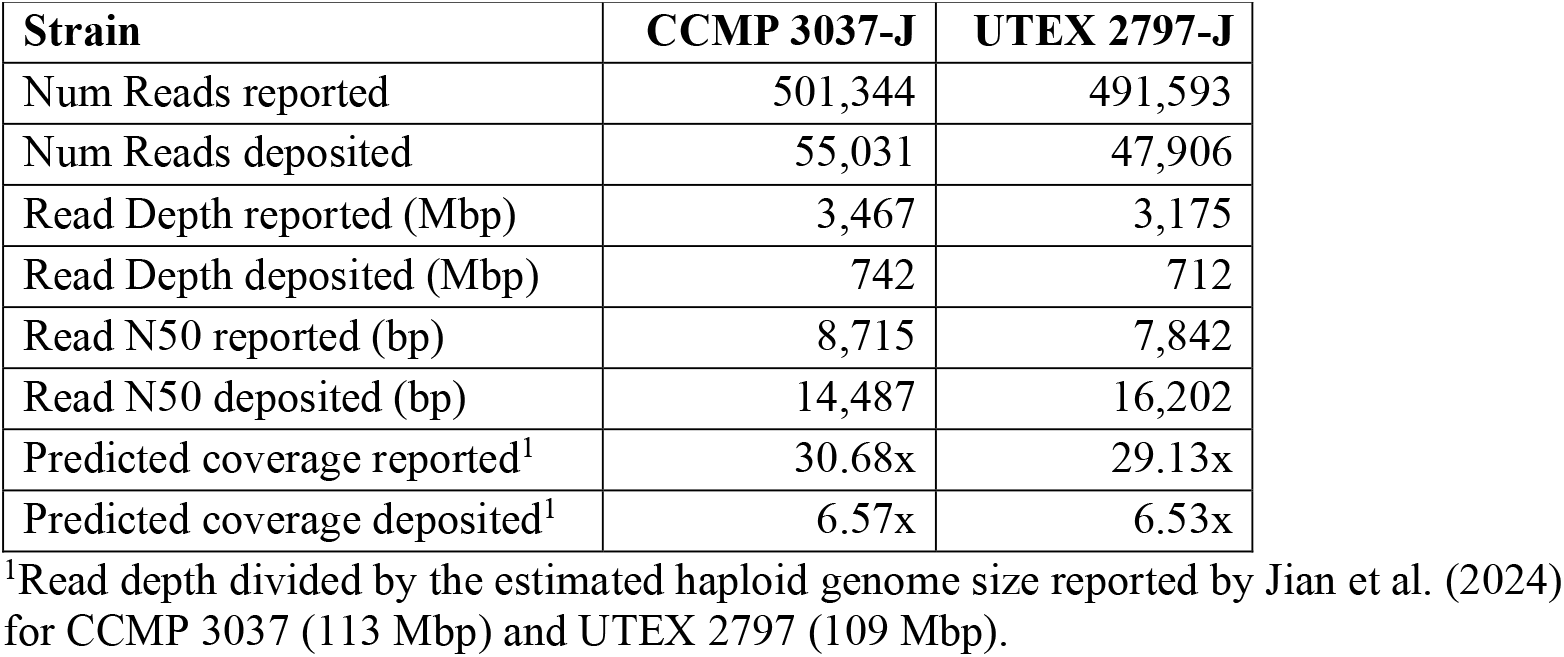
HiFi sequencing data as reported by Jian et al. (2024) compared to published dataset deposited into the CNGBdb.

| Strain | CCMP 3037-J | UTEX 2797-J |
| --- | --- | --- |
| Num Reads reported | 501,344 | 491,593 |
| Num Reads deposited | 55,031 | 47,906 |
| Read Depth reported (Mbp) | 3,467 | 3,175 |
| Read Depth deposited (Mbp) | 742 | 712 |
| Read N50 reported (bp) | 8,715 | 7,842 |
| Read N50 deposited (bp) | 14,487 | 16,202 |
| Predicted coverage reported<sup>1</sup> | 30.68x | 29.13x |
| Predicted coverage deposited<sup>1</sup> | 6.57x | 6.53x |

<sup>1</sup>Read depth divided by the estimated haploid genome size reported by Jian et al. (2024) for CCMP 3037 (113 Mbp) and UTEX 2797 (109 Mbp).

Misidentification of samples used in genomic analyses is an ongoing challenge for the field, resulting in corrections (e.g., Dixon et al. 2024) or retractions (e.g., Mu et al. 2017; Smith and Editors 2018; Christensen et al. 2021) depending on the context. The discrepancy between the two UTEX 2797 assemblies has propagated confusion and resulted in the strain’s exclusion from a subsequent genomic comparison of *P. parvum* s.l. (see Kuhl et al. 2024). This is unfortunate as UTEX 2797 has been used in numerous prior studies of the species complex (e.g., Blossom et al. 2014; Rashel and Patiño 2017; Lysgaard et al. 2018; Binzer et al. 2019; Richardson and Patiño 2021; Chávez Montes et al. 2024) and should not be omitted from genome-level investigations. The elevated heterozygosity present in UTEX 2797 is an intriguing aspect of its biology worthy of further study. Strain misidentification is a particular concern when working with microbial and cryptic species such as *P. parvum* s.l. because the lack of morphological identifiers necessitates that mix ups be detected via sequencing. In addition to correcting the record, our goal with this letter is to illustrate the dire need for clade-specific markers to efficiently confirm strain identity in *P. parvum* s.l. and microbial cryptic species complexes more generally.

## Materials and methods

### Data sources

Assemblies were downloaded from CNGB and Figshare for assemblies from Jian et al. (2024) and Wisecaver et al. (2023), respectively. Statistics were calculated using BBTools v38.91 stats.sh.

### Illumina sequencing

Fresh cultures of UTEX 2797 and CCMP 3037 were acquired directly from their respective culture collections. Genomic DNA was isolated from each strain and Illumina short-read sequencing was performed (Novogene Corporation Inc., Sacramento, CA) as described in

Wisecaver et al. (2023). Briefly, cultures were grown to late-exponential phase before pelleting and snap-freezing in liquid nitrogen. DNA was extracted from cell-pellets using a CTAB-based protocol which can be read here https://doi.org/10.17504/protocols.io.b5qhq5t6 (Auber 2019).

### Read processing and k-mer counting

UTEX 2797 and CCMP 3037 short-read libraries from both publications were compared alongside the resequenced libraries generated in this study (Wisecaver et al. 2023; Jian et al. 2024). All reads were processed according to the pipeline in Wisecaver et al. (2023). Briefly, bacterial contamination was flagged and removed using BlobTools v1.1.1 (Laetsch and Blaxter 2017), and the filtered Illumina reads were randomly subsampled to equal read counts (n = 40 million pairs) with BBMap reformat.sh v38.87 to facilitate comparison across plots. K-mers were counted with KMC v3.1.1 (Kokot et al. 2017). K-mers were also counted on unfiltered long-read sequences from both publications using KMC v3.1.1 (Kokot et al. 2017).

### SNP analysis

Two reference genomes are currently available for *P. parvum* s.l.: the prymnesin type A producing strain 12B1 sequenced by Wisecaver et al. (2023) and the type B producing strain ODER1 sequenced by Kuhl et al. (2024). To avoid reference bias that could occur when aligning to the type A genome, reads were aligned to haplotype 1 of the ODER1 type B genome (Kuhl et al. 2024) using BWA-MEM v0.7.17 (Li 2013) for Illumina short reads and Minimap2 v2.28 (Li 2018) for ONT and PacBio long reads. In addition, we downloaded Illumina RNAseq reads for UTEX 2797 sequenced by a third party (Kuhl et al. 2024), which were aligned to the ODER1 genome using STAR v2.7.11b (Dobin et al. 2013). All properly paired read alignments were filtered to remove read duplicates and secondary/chimeric alignments using Samtools v1.9 and GATK v4.2.6.1 MarkDuplicates (Li et al. 2009; Brouard et al. 2019; Van der Auwera and O’Connor 2020; Danecek et al. 2021).

SNPs were called using BCFtools v1.17 mpileup (Danecek et al. 2021) and filtered to only those present in all samples with a minimum quality of 30 using VCFtools v0.1.16x (Danecek et al. 2011), which yielded 77,168 total SNPs. For PCA independent variants (n=6,783) were extracted using PLINK v1.90b6.21 --extract with a window size of 50kb, step size of 10kb, and r^2^ of 0.1 and analyzed with PLINK v1.90b6.21 --PCA. All 77,168 SNPs were used to generate a distance matrix using PLINK v1.90b6.21 --distance square which was visualized using NeighborNet implemented in SplitsTree4 v4.19.2 with options lambda frac 1 and variance OrdinaryLeastSquares.

## Data Availability

Wisecaver et al. (2023) reads were downloaded from the Short Read Archive (SRA) of the National Center for Biotechnology Information (NCBI), BioProject PRJNA807128. Jian et al. (2024) reads were downloaded from the CNGB Sequence Archive (CNSA) of China National GeneBank DataBase (CNGBdb), accession number CNP0004284. Kuhl et al. (2024) were downloaded from the NCBI SRA, BioProject PRJNA1101072. For resequencing, strain UTEX 2797 was acquired from the UTEX Culture Collection of Algae at the University of Texas at Austin (https://utex.org) and strain CCMP 3037 was acquired from the National Center for Marine Algae and Microbiota at the Bigelow Laboratory for Ocean Sciences (https://ncma.bigelow.org). Resequenced UTEX 2797 and CCMP 3037 reads have been deposited in the NCBI SRA under accessions SRR34640331 and SRR34640332.

## Acknowledgements

We would like to thank Jo Ann Banks, Abigail Lind, and Brian Dilkes for providing helpful feedback. This research used resources of the Center for Institutional Research Computing at Washington State University and the Rosen Center for Advanced Computing at Purdue University. Funds for this work were provided by Washington State University through startup funds to JHW.

## References

Auber R. 2019. Total DNA extraction from plant tissue using CTAB method. protocols.io [Internet]. Available from: 10.17504/protocols.io.bamnic5e

Binzer SB, Svenssen DK, Daugbjerg N, Alves-de-Souza C, Pinto E, Hansen PJ, Larsen TO, Varga E. 2019. A-, B- and C-type prymnesins are clade specific compounds and chemotaxonomic markers in Prymnesium parvum. Harmful Algae 81:10–17.

Blossom HE, Rasmussen SA, Andersen NG, Larsen TO, Nielsen KF, Hansen PJ. 2014. Prymnesium parvum revisited: relationship between allelopathy, ichthyotoxicity, and chemical profiles in 5 strains. Aquat. Toxicol. Amst. Neth. 157:159–166.

Brouard J-S, Schenkel F, Marete A, Bissonnette N. 2019. The GATK joint genotyping workflow is appropriate for calling variants in RNA-seq experiments. J. Anim. Sci. Biotechnol. 10:44.

Bryant D. 2003. Neighbor-Net: An Agglomerative Method for the Construction of Phylogenetic Networks. Mol. Biol. Evol. 21:255–265.

Chávez Montes RA, Mary MA, Rashel RH, Fokar M, Herrera-Estrella L, Lopez-Arredondo D, Patiño R. 2024. Hormetic and transcriptomic responses of the toxic alga Prymnesium parvum to glyphosate. Sci. Total Environ. 954:176451.

Cheng H, Concepcion GT, Feng X, Zhang H, Li H. 2021. Haplotype-resolved de novo assembly using phased assembly graphs with hifiasm. Nat. Methods 18:170–175.

Christensen KA, Rondeau EB, Minkley DR, Leong JS, Nugent CM, Danzmann RG, Ferguson MM, Stadnik A, Devlin RH, Muzzerall R, et al. 2021. Retraction: The Arctic charr (Salvelinus alpinus) genome and transcriptome assembly. PLOS ONE 16:e0247083.

Danecek P, Auton A, Abecasis G, Albers CA, Banks E, DePristo MA, Handsaker RE, Lunter G, Marth GT, Sherry ST, et al. 2011. The variant call format and VCFtools. Bioinformatics 27:2156–2158.

Danecek P, Bonfield JK, Liddle J, Marshall J, Ohan V, Pollard MO, Whitwham A, Keane T, McCarthy SA, Davies RM, et al. 2021. Twelve years of SAMtools and BCFtools. GigaScience 10:giab008.

Dixon OFL, De Silva C, Phillips BT, Gallagher AJ. 2024. Correction to: Novel deep-sea observations reveal twilight zone occurrence for two species of pelagic sharks: the bignose shark Carcharhinus altimus and the silky shark Carcharhinus falciformis. Environ. Biol. Fishes 107:1171–1174.

Dobin A, Davis CA, Schlesinger F, Drenkow J, Zaleski C, Jha S, Batut P, Chaisson M, Gingeras TR. 2013. STAR: ultrafast universal RNA-seq aligner. Bioinformatics 29:15–21.

Huson DH. 1998. SplitsTree: analyzing and visualizing evolutionary data. Bioinformatics 14:68– 73.

Jian J, Wu Z, Silva-Núñez A, Li X, Zheng X, Luo B, Liu Y, Fang X, Workman CT, Larsen TO, et al. 2024. Long-read genome sequencing provides novel insights into the harmful algal bloom species Prymnesium parvum. Sci. Total Environ. 908:168042.

John U, Litaker RW, Montresor M, Murray S, Brosnahan ML, Anderson DM. 2014. Formal revision of the Alexandrium tamarense species complex (Dinophyceae) taxonomy: The introduction of five species with emphasis on molecular-based (rDNA) classification. Protist 165:779–804.

Kokot M, Długosz M, Deorowicz S. 2017. KMC 3: counting and manipulating k-mer statistics. Bioinformatics 33:2759–2761.

Kuhl H, Strassert JFH, Čertnerová D, Varga E, Kreuz E, Lamatsch DK, Wuertz S, Köhler J, Monaghan MT, Stöck M. 2024. The haplotype-resolved Prymnesium parvum (type B) microalga genome reveals the genetic basis of its fish-killing toxins. Curr. Biol.:S0960982224008170.

Laetsch DR, Blaxter ML. 2017. BlobTools: Interrogation of genome assemblies. F1000Research 6:1287.

Li H. 2013. Aligning sequence reads, clone sequences and assembly contigs with BWA-MEM. Available from: http://arxiv.org/abs/1303.3997

Li H. 2018. Minimap2: pairwise alignment for nucleotide sequences.Birol I, editor. Bioinformatics 34:3094–3100.

Li H, Handsaker B, Wysoker A, Fennell T, Ruan J, Homer N, Marth G, Abecasis G, Durbin R, 1000 Genome Project Data Processing Subgroup. 2009. The sequence alignment/map format and SAMtools. Bioinforma. Oxf. Engl. 25:2078–2079.

Lysgaard ML, Eckford-Soper L, Daugbjerg N. 2018. Growth rates of three geographically separated strains of the ichthyotoxic Prymnesium parvum (Prymnesiophyceae) in response to six different pH levels. Estuar. Coast. Shelf Sci. 204:98–102.

Mu X, Yang Y, Liu Y, Luo D, Xu M, Wei H, Gu D, Song H, Hu Y. 2017. Retraction Note to: The complete mitochondrial genomes of two freshwater snails provide new protein-coding gene rearrangement models and phylogenetic implications. Parasit. Vectors 10:350.

Purcell S, Neale B, Todd-Brown K, Thomas L, Ferreira MAR, Bender D, Maller J, Sklar P, De Bakker PIW, Daly MJ, et al. 2007. PLINK: A Tool Set for Whole-Genome Association and Population-Based Linkage Analyses. Am. J. Hum. Genet. 81:559–575.

Rashel RH, Patiño R. 2017. Influence of genetic background, salinity, and inoculum size on growth of the ichthyotoxic golden alga (Prymnesium parvum). Harmful Algae 66:97– 104.

Richardson ET, Patiño R. 2021. Growth of the harmful alga, Prymnesium parvum (Prymnesiophyceae), after gradual and abrupt increases in salinity. J. Phycol. 57:1335– 1344.

Simon EM, Nanney DL, Doerder FP. 2008. The “Tetrahymena pyriformis” complex of cryptic species. Biodivers. Conserv. 17:365–380.

Smith CM, Editors TPO. 2018. Retraction: Virulent Diuraphis noxia Aphids Over-Express Calcium Signaling Proteins to Overcome Defenses of Aphid-Resistant Wheat Plants. PLOS ONE 13:e0191678.

Sonneborn TM. 1975. The Paramecium aurelia complex of fourteen sibling species. Trans. Am. Microsc. Soc. 94:155–178.

Van der Auwera GA, O’Connor BD. 2020. Genomics in the Cloud: Using Docker, GATK, and WDL in Terra. O’Reilly Media

Wisecaver JH, Auber RP, Watervoort NF, Fallon TR, Riedling OL, Manning SR, Moore BS, Driscoll WW. 2023. Extreme genome diversity and cryptic speciation in a harmful algal-bloom-forming eukaryote. Curr. Biol. 33:2246-2259.e8.

